# Active Poroelastic Two-Phase Model for the Motion of Physarum Microplasmodia

**DOI:** 10.1101/638312

**Authors:** Dirk Alexander Kulawiak, Jakob Löber, Markus Bär, Harald Engel

**Affiliations:** TU Berlin - Institut für Theoretische Physik, Hardenberstr. 36, 10623 Berlin, Germany; Max-Planck-Institut für Physik komplexer Systeme, Nöthnitzer Straße 38, 01187 Dresden, Germany; Physikalisch-Technische Bundesanstalt, Abbestrasse 2-12, 10587 Berlin, Germany

## Abstract

The onset of self-organized motion is studied in a poroelastic two-phase model with free boundaries for Physarum microplasmodia (MP). In the model, an active gel phase is assumed to be interpenetrated by a passive fluid phase on small length scales. A feedback loop between calcium kinetics, mechanical deformations, and induced fluid flow gives rise to pattern formation and the establishment of an axis of polarity. Altogether, we find that the calcium kinetics that breaks the conservation of the total calcium concentration in the model and a nonlinear friction between MP and substrate are both necessary ingredients to obtain an oscillatory movement with net motion of the MP. By numerical simulations in one spatial dimension, we find two different types of oscillations with net motion as well as modes with time-periodic or irregular switching of the axis of polarity. The more frequent type of net motion is characterized by mechano-chemical waves traveling from the front towards the rear. The second type is characterized by mechano-chemical waves that appear alternating from the front and the back. While both types exhibit oscillatory forward and backward movement with net motion in each cycle, the trajectory and gel flow pattern of the second type are also similar to recent experimental measurements of peristaltic MP motion. We found moving MPs in extended regions of experimentally accessible parameters, such as length, period and substrate friction strength. Simulations of the model show that the net speed increases with the length, provided that MPs are longer than a critical length of *≈* 120 µm. Both predictions are in line with recent experimental observations.

## 1 Introduction

Dynamic processes in biological systems such as cells are examples of when spatio-temporal patterns develop far from thermodynamic equilibrium [1, 2]. One fascinating instance of such active matter are intracellular molecular motors that consume ATP [3] and can drive mechano-chemical contraction-expansion patterns [4] and, ultimately, cell locomotion. Further biological examples of such phenomena are discussed in [5–7].

The true slime mold Physarum polycephalum is a well known model organism [8] that exhibits mechano-chemical spatio-temporal patterns. Previous research in Physarum has addressed many different topics in biophysics, such as genetic activity [9], habituation [10], decision making [11] and cell locomotion [12, 13]. Physarum is an unicellular organism, which builds large networks that exhibit self-organized synchronized contraction patterns [8, 14, 15]. These contractions enable shuttle streaming in the tubular veins of the network and allow for efficient nutrient transport throughout the organism [16]. Many groups have investigated the network’s dynamics [17–19], however size and complex topology of these networks make analyzing and modeling them challenging.

Physarum microplasmodia (MP) allow one to study Physarum’s internal dynamics in a simpler setup. These MPs can be produced by extracting cytoplasm from a vein and placing it on a substrate. After reorganization, these droplets of cytoplasm show a surprising wealth of spatio-temporal patterns such as spiral, standing and traveling waves and irregular and anti-phase oscillations in their height [20]. After several hours, MPs are deforming to a tadpole-like shape and start exploring their surroundings [21–24].

MP motion is composed of an oscillatory forward and backward motion. Their forward motion has a larger magnitude than the backward motion. There are experimental observations of two distinct motility types: *peristaltic* and *amphistaltic* [13, 24]. In the more common peristaltic case forward traveling contraction waves result in forward motion of the whole cell body. In the amphistaltic type, standing waves cause front and rear to contract in anti-phase.

Different modes of motion driven by periodic deformation waves are not restricted to Physarum MP, but are also found in rather general mathematical models [25] and in the locomotion of a wide variety of limbless and legged animals [26]. More recently, different modes have also been found in the Belousov-Zhabotinsky reaction in a light-driven photosensitive gel [27–30]; Epstein and coworkers have classified the modes using the terms *retrograde* and *direct wave locomotion* [28] and also report on a form of oscillatory migration without net displacement of the average position [29].

Common models for cell locomotion and the cytoskeleton’s dynamics are based on active fluid and gel models [31–34]. While simple fluids and solids are governed by a single momentum balance equation, poroelastic media belong to the class of two-fluid models and possess individual momentum balance equations for each of the phases. This approach is useful if the two constituent phases have largely different rheological properties and penetrate each other on the relatively small length scales on which cytosol permeates the cytoskeleton [35].

To describe the MP’s motion, we utilize an active poroelastic two-phase description [35, 36]. Active poroelastic models have been used to describe the pattern formation in resting [37] and moving [38] poroelastic droplets. In simple generic models, a feedback loop between a chemical regulator, mechanical contractions and induced flows give rise to pattern formation in a resting droplet [37]. In [38], we have studied this model with free boundary conditions and linear friction between droplet and substrate. While we were able to observe back and forth motion of the boundaries, the center of mass (COM) position remained fixed. The droplet did not exhibit net motion. Moreover, we derived an argument that COM motion is impossible with a spatially homogeneous substrate friction. Furthermore, numerical simulations have shown that an additional mechanism establishing an axis of polarity that is stable on long timescales is necessary for net motion [38].

In this work, we proceed in two steps. First we extend the model described in [38] with a nonlinear slip-stick friction between droplet and substrate to create a spatially heterogeneous substrate friction. Second, we consider a specific model derived for Physarum MP that contains a reaction kinetics. Within this enhanced model, we explore the conditions for the onset of motion of Physarum MP.

Previous work on models for resting Physarum MP by Radszuweit et al. has shown that inclusion of a nonlinear reaction kinetics for the calcium regulator can result in the emergence of uni-directional traveling mechano-chemical waves that establish an axis of polarity inside the MP [39, 40]. Nonlinear friction is a common assumption in biology [41, 42] and can also be found in other eucaryotic cells [43, 44]. Here, we utilize a simplified version of the nonlinear slip-stick friction model introduced by Barnhart et al. to account for oscillatory modulations of keratocyte movement [44]. The nonlinear friction dynamics and its synchronization with other properties such as the local strain inside a cell are also important for other types of amoeboid motion such as chimneying [45]. Moreover, regulation of the friction strength is significant for locomotion in other biological systems such as snails or slugs [46, 47].

Section 2 contains a comprehensive description of the model. In Section 3, we analyze the linear stability of the model and identify different types of motion. Furthermore, we explore the effects of parameters that are accessible in the experiment such as the period of the internal dynamics, the length and the substrate friction strength. Section 3 contains a comparison of our findings with recent experiments of Physarum MP. Afterwards, we summarize our results in the discussion and briefly address possible extensions of the model.

## 2 Model

We follow our earlier work [37–40, 48, 49] and utilize an active poroelastic two-phase model in one spatial dimension to describe homogeneous and isotropic Physarum microplasmodia (MP). We assume that MP consist of an active gel phase representing the cytoskeleton that we model as a viscoelastic solid with a displacement field *u* and velocity field 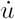 [50]. The gel is penetrated by a passive cytosolic fluid phase with flow velocity field *v*. For related models see [51, 52].

The total stress in the medium is given by *σ* = *ρ*_*g*_*σ*_*g*_ + *ρ*_*f*_ *σ*_*f*_, where *ρ*_*g*_ (*ρ*_*f*_) denotes the volume fraction and *σ*_*g*_ (*σ*_*f*_) the stress tensor in gel and fluid phase, respectively. While the time evolution for the gel fraction is given by 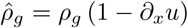, only the constant term *ρ*_*g*_ enters into the equations due to our small strain approximation with | *∂*_*x*_*u* | ≪ 1. Assuming that no other phases are present, the volume fractions obey *ρ*_*g*_ + *ρ*_*f*_ = 1 [53].

Each phase satisfies a momentum balance equation *∂*_*x*_(*σ*_*g/f*_−*p*) + *f*_*g/f*_ + *f*_sub_ = 0, where *p* denotes the hydrodynamic pressure. The friction between both phases is given by Darcy’s law with 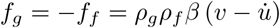 Following [44], we assume that the friction between gel and substrate depends nonlinearly on the the local gel speed according 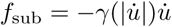 and no friction between fluid and substrate.

We model MPs in an one-dimensional time-dependent domain *𝓑* with boundaries denoted by *∂𝓑*. Assuming that *𝓑* is infinitely large in the y-direction, the boundary is straight, and we omit terms that depend on interface tension or bending. Free boundary conditions in x-direction enable boundary deformations and thus motion in response to bulk deformations [54–56]. The total stress has to be continuous across the boundary and with the assumption that the MP is embedded in an inviscid fluid, the first boundary condition is given by

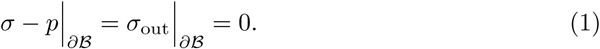

The model is composed of two phases with individual momentum balances. Hence, we need a second boundary condition to close the model equations. Neglecting permeation of gel or fluid through the boundary, the second boundary condition is

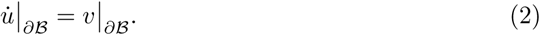

Free boundary conditions require solving the momentum balances at the boundary, whose position must be determined in the course of solving the evolution equations. To circumvent this problem, we formulate the model in a co-moving body reference frame. The details of the transformation from the laboratory to the body reference frame can be found in [37–39]. S1 Fig displays a visual comparison of a quantity plotted in the body reference and the laboratory frame.

The stress of the passive fluid with viscosity *η*_*f*_ is given by *σ*_*f*_ = *η*_*f*_ *∂*_*x*_*v*. The gel’s stress can be decomposed in a passive part, *σ*^ve^, and an active part, *σ*^act^. The passive part is described as a viscoelastic Kelvin-Voigt solid with 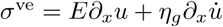[50,57] where *E* denotes Young’s modulus and *η*_*g*_ the dynamic viscosity. For a discussion on the effects of different linear and nonlinear viscoelastic models see [36, 58]. The active stress is assumed to be governed by a chemical regulator *c*, which is usually identified as calcium [13, 59, 60], according to

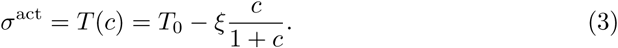

In this formulation, the homogeneous part of the active stress *T*_0_ is inhibited by calcium concentration *c* with a coupling strength *ξ* > 0.

In [38], we identified a heterogeneous substrate friction as a requisite for motion of the MP’s center of mass (COM). While the exact nature of the Physarum friction dynamics is not known, recent experiments indicate that MP exhibit nonlinear slip-stick friction with an underlying substrate resulting in a heterogeneous friction [13]. We assume that there are adhesive bonds between gel and substrate which break once the force acting on them is larger than a critical value and that this force depends on the local gel speed 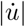 [44]. Therefore, if 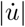 is larger than the critical slip speed *v*_slip_ these bonds will break and the friction coefficient will decrease locally, yielding

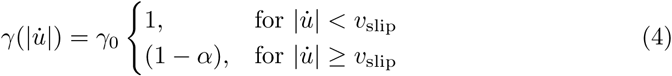

with 0 < *α* < 1. With *α* = 0 the friction coefficient is always homogeneous, higher values allow for heterogeneous friction coefficient values. In the following, we call *α* the slip-ratio and *γ*_0_ the base friction coefficient.

The fluid phase contains dissolved chemical species that can perform regulatory activities. Following [40], we consider an advection-diffusion-reaction dynamics for two control species with local instantaneous concentrations: calcium *c* and a control species *a* that chemically interacts with calcium. Calcium determines the gel’s active tension while the control species represents all biochemical interactions between calcium and other components of the cytosol. We assume an oscillatory Brusselator-type kinetics for *a* and *c* [40].

Both species are dissolved in the fluid and advected with its flow. Furthermore, they diffuse with coefficient *D*_*c*_ and *D*_*a*_, respectively. We assume that both species cannot cross the boundary, resulting in no-flux boundary conditions.

Linearizing with respect to the gel strains *∂*_*x*_*u* yields advection-diffusion-reaction equations with the relative velocity of the fluid to the gel 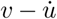 as the advection velocity. In the body reference frame the equations for *c* and *a* read

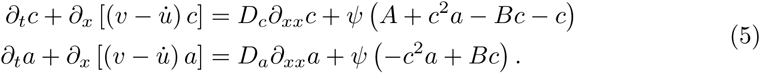

Here, *ψ* = 0.105/s represents the temporal scale of the calcium kinetics which can be used to fit the experimentally observed time scale of the mechano-chemical waves.

The homogeneous steady state (HSS) of the model is given by *c* = *A* and *a* = *B/A*. The HSS destabilizes through a Hopf-Bifurcation if *B* > *B*_cr_ = 1 + *A*^2^ [40]. Here, we fix *A* = 0.8 and then vary *B* above its critical value *B*_cr_. With the inclusion of a calcium kinetics, the total amount of calcium is not conserved anymore. In addition, the calcium kinetics can introduce an axis of polarity [39, 40]. The case with calcium conservation was examined in earlier studies for resting [37] and moving poroelastic droplets [38].

Fig. 1 gives an overview of the different elements in the model. Calcium regulates the gel’s activity. Spatial variations in the calcium concentration yield a heterogeneous active tension which results in deformation and motion of the gel. Once the gel moves, there is nonlinear friction between gel and substrate. Furthermore, the gel’s motion induces pressure gradients. Due to these gradients and friction between the gel and the fluid phase, the fluid starts to flow and the dissolved chemical species are advected with the fluid, closing the feedback loop between the different elements in the model.

**Fig 1.**
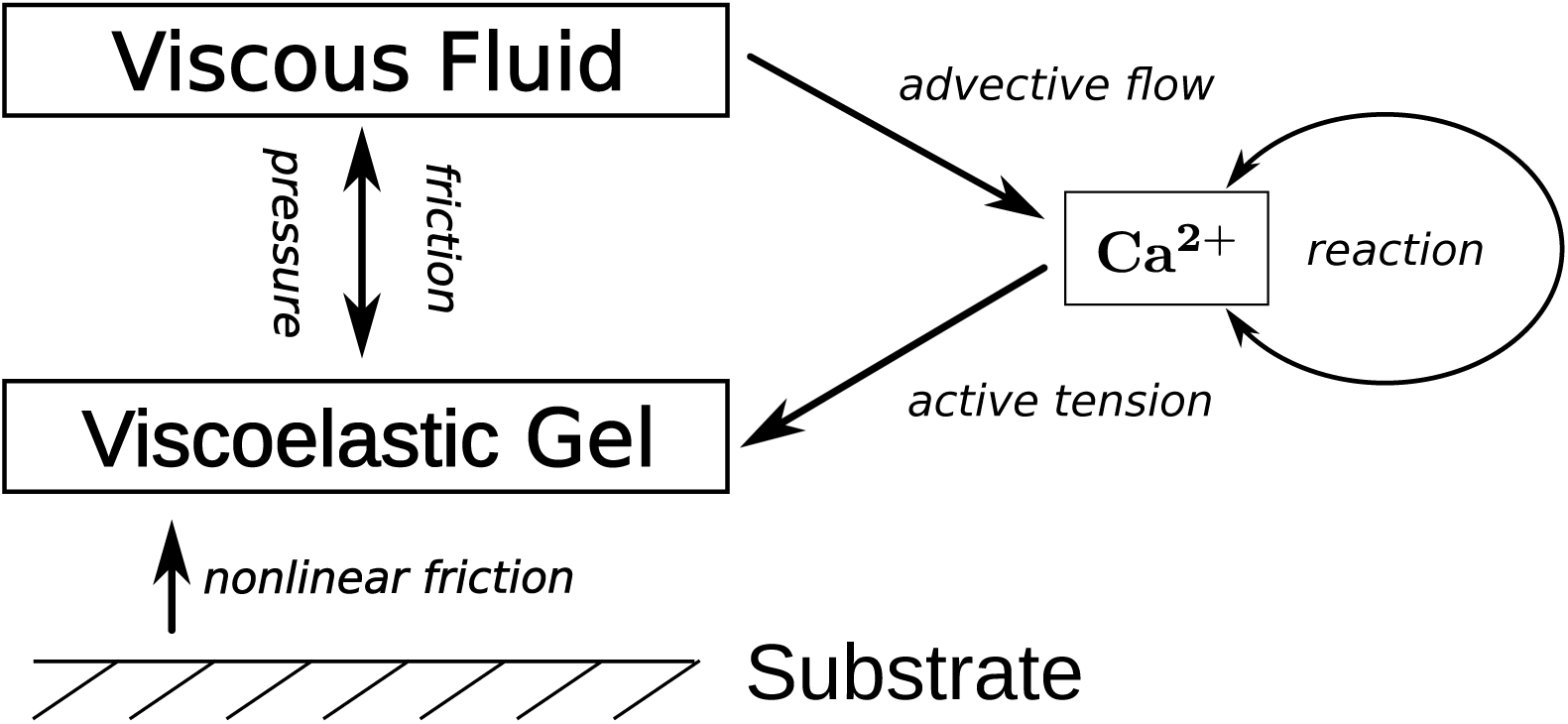
Sketch of the model. Calcium regulates the gel’s active tension and varying calcium concentration levels cause gel deformations. When the gel moves, there is nonlinear friction between gel and substrate. Moreover, pressure gradients arise and there is friction between both phases. Both cause the fluid to start flowing and advecting the dissolved chemical species.

In summary, the model equations are given by

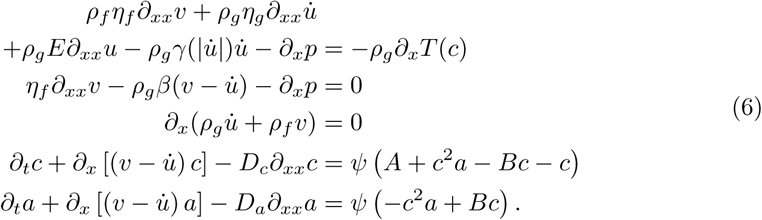

Following [39], we introduce the Péclet number Pe = *ξ*/(*D*_*c*_*β*) as a measure for the ratio of diffusive to advective time scales to characterize the strength of the active tension [37]. We also define the dimension-less ratio of the Péclet number to the critical Péclet number for the onset of the mechano-chemical instability without a calcium kinetics [40] as *F* = *ξA*/ (*D*_*c*_*β* (1 + *A*)^2^). For *F* > 1(*F* < 1), the HSS is stable (unstable).

## 3 Results

We solve Eq. (6) numerically in a one-dimensional domain of length *L* with parameters adopted from [40] which are listed in S1 Table, unless stated otherwise. Our initial condition is the weakly perturbed homogeneous steady state (HSS) with

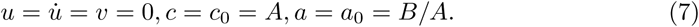

The length *L* remains constant due to the incompressibility of the medium. We use the center of mass (COM) position 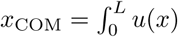 as a measurement of the position. Note that we will compare the COM’s motion to our results from [38] where we utilized the location of the left boundary as the position and the COM always remained fixed. In addition, we utilize the temporal distance between subsequent peaks in the calcium concentration *c* as a measure for the period *P* of the internal dynamics.

Depending on the parameter values of active tension (F), nonlinear friction (*α, v*_slip_, *γ*_0_) and calcium kinetics (*B, ψ*), we can observe resting, stationary MPs and MPs performing three types of oscillatory motion: first, with fixed COM; second, with back and forth moving COM; third, with back and forth moving COM together with net displacement of the COM (net motion). In addition, we analyze the effect of parameters that are accessible in the experiment like the length *L* and the base friction coefficient *γ*_0_.

### Nonlinear slip-stick friction allows for COM motion

First, we analyze the effect of the nonlinear substrate friction without calcium kinetics (*ψ* = 0), i. e. the calcium dynamics is governed by a pure advection-diffusion dynamics and the total amount of calcium is conserved. Afterwards, we introduce a nonlinear calcium kinetics to establish an axis of polarity.

We vary the slip-ratio *α* and the slip-velocity *v*_slip_. For spatially uniform substrate friction (*α* = 0), there are gel deformations, and the boundaries change their position, but the COM’s position remains fixed as described in [38]. Increasing *α* > 0 gives rise to spatially heterogeneous friction allowing for COM motion. For small increases of *α*, the mode of motion usually remains the same as with a homogeneous friction, but it is now accompanied by COM motion of the same type but with a smaller magnitude. In general, the COM motion’s amplitude will increase when increasing *α* (S2 Fig). However, large values of *α* might result in a change of the mode of motion, for example, an initially periodic dynamics might become irregular. For values of 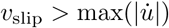 the friction is always in the sticking regime. Therefore, the friction coefficient is homogeneous. Likewise, for low values of *v*_slip_ the moving MP is almost always in the regime of slipping friction resulting in an effectively homogeneous friction coefficient of *γ* = *γ*_0_(1 *-α*). Intermediate values of *v*_slip_ result in a heterogeneous distribution of the friction coefficient.

Comparable to our results in [38], the position over time remains fixed or undergoes periodic (irregular) motion of the COM and the boundaries together with a periodic (irregular) calcium dynamics, as shown in Fig. 2. However, we found no cases of net motion. As discussed in [38], the lack of net motion gives rise to the question of why the time-averaged position vanishes for all of these cases. While the calcium distributions produced by pure advection-diffusion dynamics may look asymmetric at certain instances in time, the long time-averaged distribution is always symmetric. This indicates that the front-back symmetry is not broken on timescales of *t* ≫ *P*, and that an additional mechanism to establish an axis of polarity is needed.

**Fig 2.**
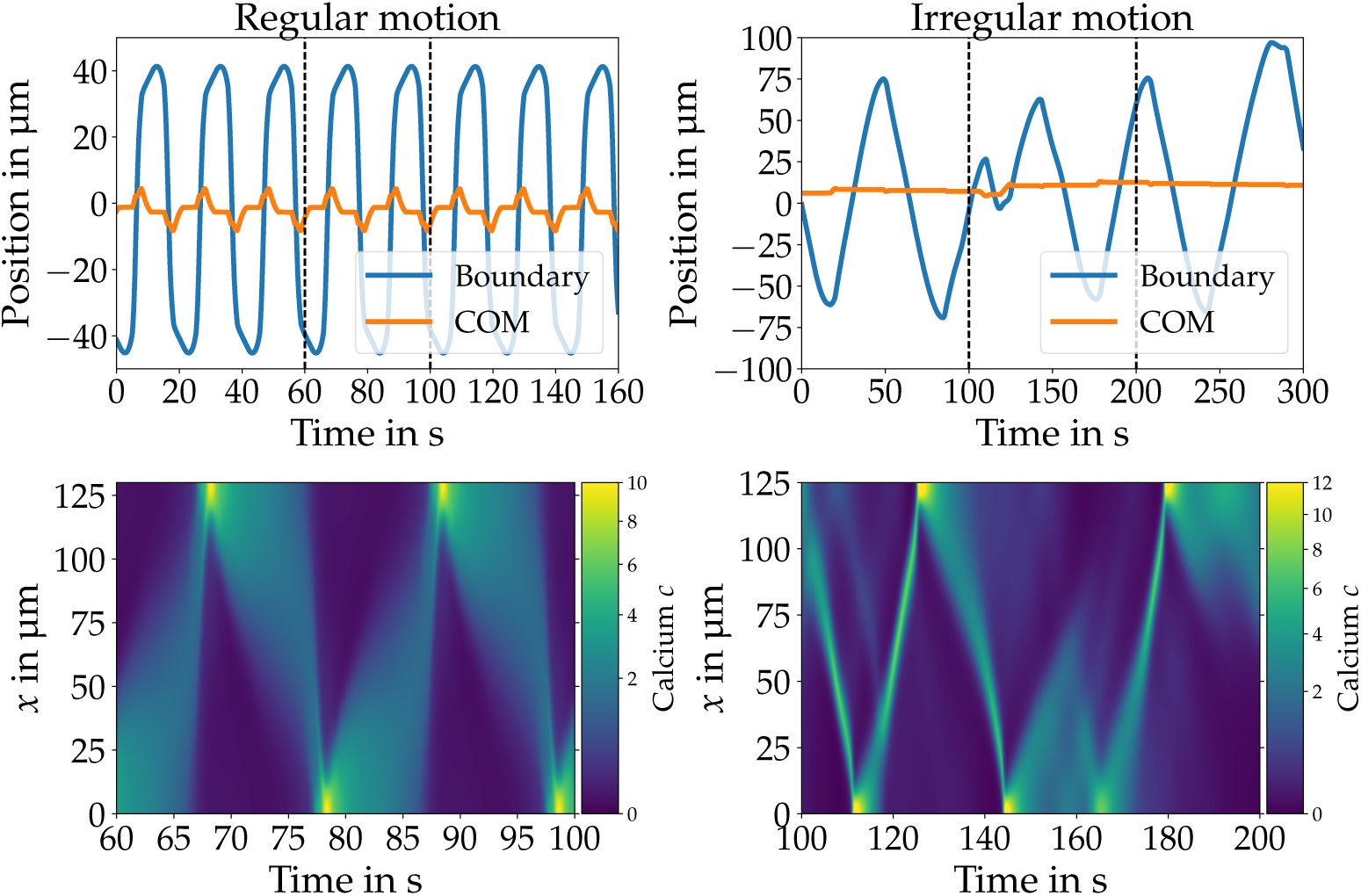
COM (orange) and boundary (blue) trajectories (top row) and calcium dynamics (bottom row) for regular (left column) and irregular (right column) motion with nonlinear substrate friction without calcium kinetics (*ψ* = 0). The nonlinear substrate friction creates a spatially heterogeneous friction which allows for COM motion (orange line) together with motion of the boundaries (blue). The regular calcium dynamics (bottom left) results in oscillatory back and forth motion of COM as well as the boundaries (top left). An irregular calcium dynamics (bottom right) creates irregular motion (top right). There is no net motion for both cases. Parameters: *v*_slip_ = 6.0 µm/s, *α* = 0.25, *γ*_0_ = 10^−5^ kg/s, *L* = 130 µm, Pe = 50 (left) and *v*_slip_ = 30.0 µm/s, *α* = 0.3, *γ*_0_ = 10^−5^ kg/s, Pe = 9.5, *β* = 10^−4^ kg µm^−3^ s^−1^, *E* = 0.01kg µm^−1^ s^−2^, *η*_*g*_ = 0.01kg µm^−1^ s^−1^, *η*_*f*_ = 2 × 10^−8^ kg µm^−1^ s^−1^ (right).

Note that the exact nature of Physarum’s friction dynamics can not be inferred from the available experimental data. A nonlinear dependency of *γ* on other quantities such as the calcium concentration *c*, the local contraction amplitude *∂*_*x*_*u* (which is proportional to the height *h* [39]), and more complex slip-stick models led to qualitatively equivalent results for the onset of COM motion.

### Nonlinear calcium kinetics creates an axis of polarity

One self-organized way that leads to the emergence of uni-directional traveling mechano-chemical waves which establishes an axis of polarity that is stable on timescales of *t* ≫ *P* is to introduce a nonlinear calcium kinetics as studied in [39, 40]. Introducing this calcium kinetics allows for a temporal variation of the total amount of calcium. The amplitude of calcium waves can vary while traveling waves can annihilate on collision with a boundary. This is in contrast to the behavior with a pure advection-diffusion dynamics where calcium is conserved and waves always get reflected at the boundaries.

The linear stability of the HSS (Eq. 7) against small perturbations is analyzed in detail in [39, 40]. In [40] five regions with qualitatively different dispersion relations are given: i) the HSS is stable for small *F* and *B*. ii) Upon increasing the chemical activity *B* > *B*_cr_ the HSS undergoes a supercritical Hopf bifurcation. iii) Decreasing *B* < *B*_cr_ and increasing the mechanical activity *F* > 1 gives rise to an oscillatory short-wavelength instability as already identified and discussed in [40]. iv) For *B* > *B*_cr_ and *F* > 1 both instabilities are present and more complex wave patterns can emerge. v) A further increase of *F* results in the real eigenvalues maximum to become purely real for small *k*. However, for larger *k* these eigenvalues still have an imaginary component ≠ 0. Note that we analyzed the linear stability of the model with linear friction 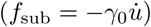, as the nonlinear friction terms vanishes in the linear approximation. The forms of the typical resulting dispersion relations are analogous to the ones displayed in [40].

In numerical simulations, we find modes defining an axis of polarity that is stable on timescales of *t* ≫ *P* for parameters in region iv. These modes are distinguished by a spatially asymmetric long time-averaged calcium distribution. This is in contrast to the averaged calcium distributions produced by the pure advection-diffusion dynamics which are always spatially symmetric. Such an axis of polarity that remains stable on long timescales will be referred to as stable polarity in the remainder of this work.

### Nonlinear calcium kinetics combined with nonlinear substrate friction result in net motion

We identified the two ingredients we need to build a model with net motion. For appropriate values for the parameters of the calcium kinetics and the substrate friction oscillatory back and forth motion with net motion of the COM is observed. In the following, we characterize two types of oscillations with net motion occurring in two different parameter regions.

In the first type (type 1), calcium waves emerge periodically at the front traveling backwards with decreasing amplitude together with backward motion of both boundaries. While traveling backward, the calcium wave causes forward motion of gel and COM. Upon collision with the rear boundary, the calcium concentration first peaks and the wave is then annihilated. Once the calcium wave has decayed, the tension in the material relaxes and there is no motion until the next calcium wave arises. The slip velocity *v*_slip_ is only exceeded while a wave is propagating. Therefore, there is only COM motion during wave propagation, and the COM remains at a constant position otherwise.

The continuous emergence of calcium waves at the front and their annihilation at the back together with the asymmetric time-averaged calcium distribution indicate the establishment of a stable polarity. This dynamics results in oscillatory back and forth motion of COM and boundaries together with net motion.

The period of this motion is *P* ≈ 95 s. In each period the COM moves forward by ≈ 18 µm which results in a net speed of 0.2 µm/s. Fig. 3 shows gel velocity, calcium dynamics as well as boundary (blue) and COM (orange) position for the first type of oscillations with net motion. This type is the most commonly observed case of COM oscillations with net motion, and we observe it for a wide range of parameters.

**Fig 3.**
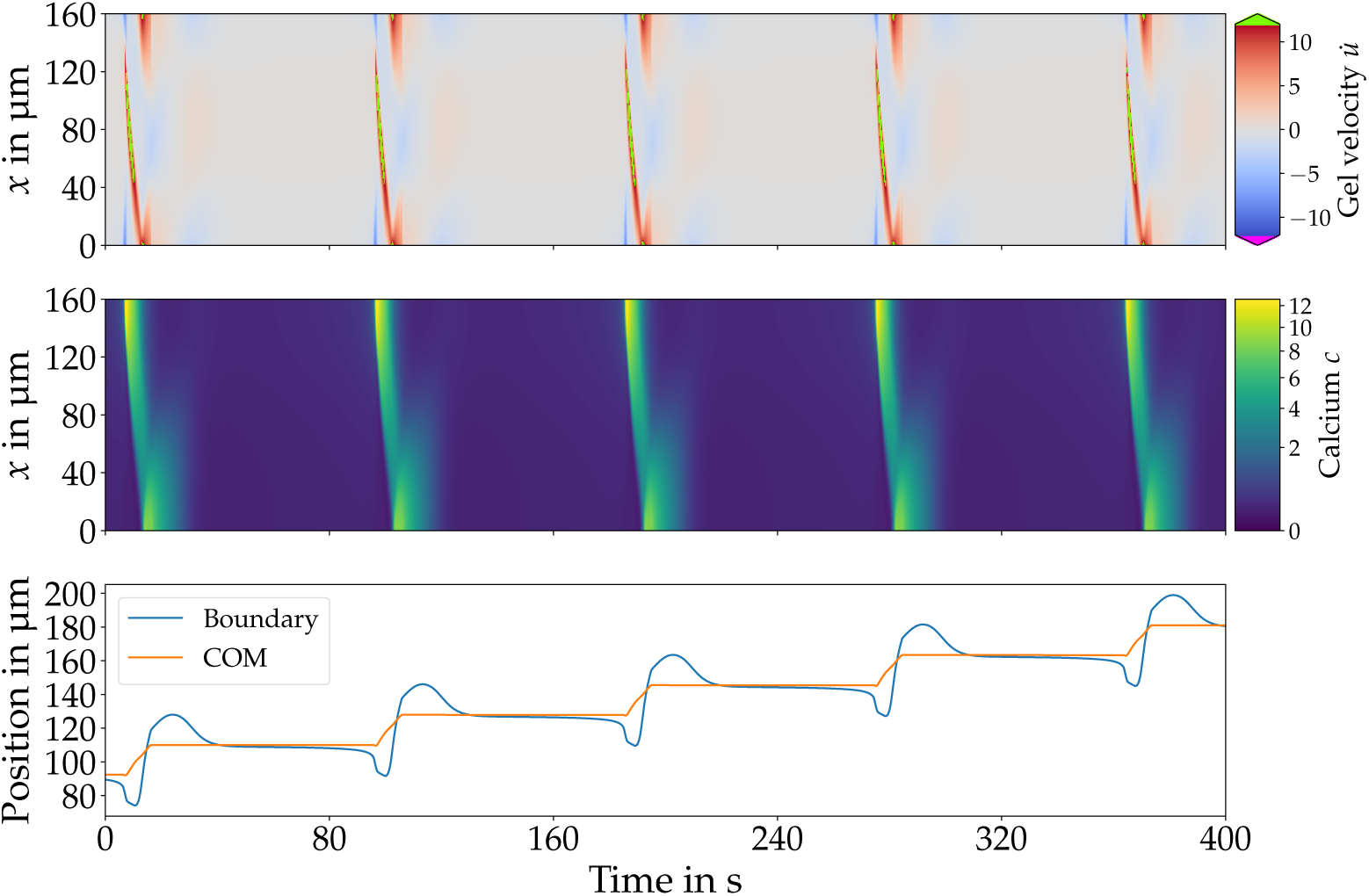
Gel velocity (top), calcium dynamics (middle) and trajectories (bottom) of boundary (blue) and COM (orange) for type 1 motion. Backward traveling calcium waves emerge periodically at the front, together with backward motion of the boundaries. Once the calcium wave is traveling backward, it causes forward gel flow and COM motion. On their arrival at the rear, the calcium concentration peaks, but the wave is annihilated upon collision with the boundary. The resulting forward gel motion is of a greater amplitude than the backward motion resulting in net motion with *v*_net_ = 0.2 µm/s. Parameters *B* = 3.5, *γ*_0_ = 7 × 10^−6^ kg/s, *v*_slip_ = 2.5 µm/s, *α* = 0.15, *L* = 160 µm and *F* = 12.3.

For the second type (type 2) of COM oscillations with net motion, calcium waves emerge alternating from both sides traveling towards the opposite boundary. Upon approaching the front (rear), the calcium waves coincide with forward (backward) motion of the boundaries. When the wave hits the boundary, the calcium concentration peaks, sparking the emergence of a new wave traveling into the opposite direction. Waves emerging from the front have a higher amplitude than waves from the back as displayed in S3 Fig. The higher amplitude of waves traveling from front to rear indicate a stable polarity and results in net motion.

The boundaries as well as the COM oscillate with the same period of *P* ≈ 14 s, however the amplitude of the COM motion is smaller than the amplitude of the boundaries motion. The COM advances the boundaries by a quarter period. In each cycle, the COM moves forward by ≈ 3 µm yielding a net speed of *v*_net_ = 0.21 µm/s. Fig. 4 shows gel velocity, calcium dynamics and COM and boundary trajectories for type 2 motion. The gel velocity resembles the pattern found in [38], but now the calcium kinetics creates a stable polarity.

**Fig 4.**
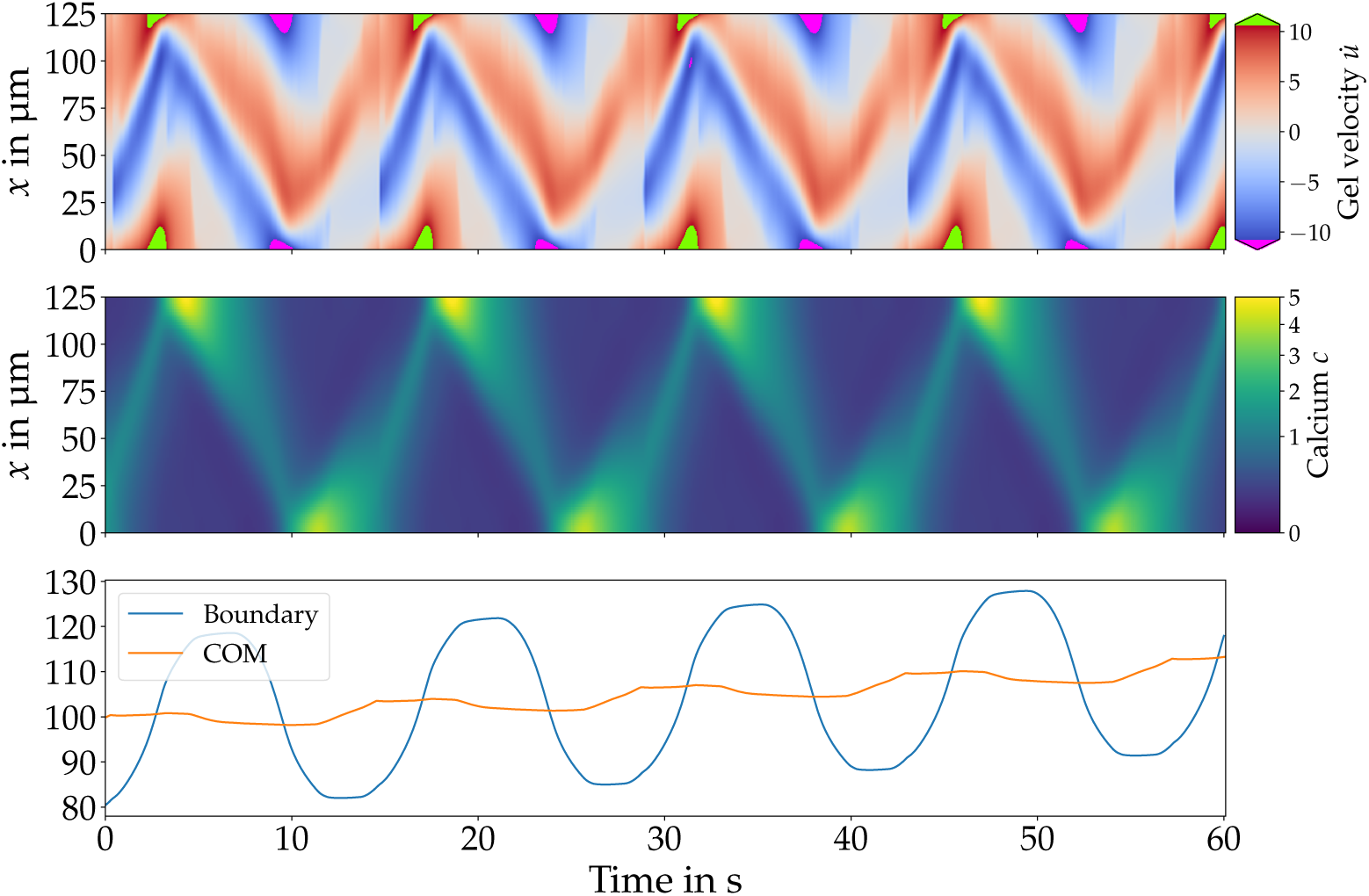
Gel velocity (left), calcium dynamics (middle) and trajectories (bottom) of boundary (blue) and COM (orange) for type 2 motion. Calcium waves emerge alternating from front and back traveling towards the opposite side. The collision with a boundary sparks a new wave traveling backwards. Waves emerging at the front have a larger amplitude than waves originating at the rear resulting in net motion. Parameters: *B* = 2, *γ*_0_ = 10^−5^ kg/s, *v*_slip_ = 1.54 µm/s, *α* = 0.1, *L* = 125 µm and *F* = 18.5.

In addition to the two types of oscillations with net motion described above, we find several modes with time-periodic or irregular switching polarity (S4 Fig).

### Model reproduces experimentally observed types of movement

Comparing our simulations with Physarum experiments reveals several qualitative and quantitative similarities. In experiments, MPs explore their surrounding with an oscillatory forward and backward motion [21, 24]. Their forward motion is of larger magnitude than their backward motion, resulting in net motion in each cycle. An experimental MP trajectory is shown in [24] (Fig 6b).

**Fig 5.**
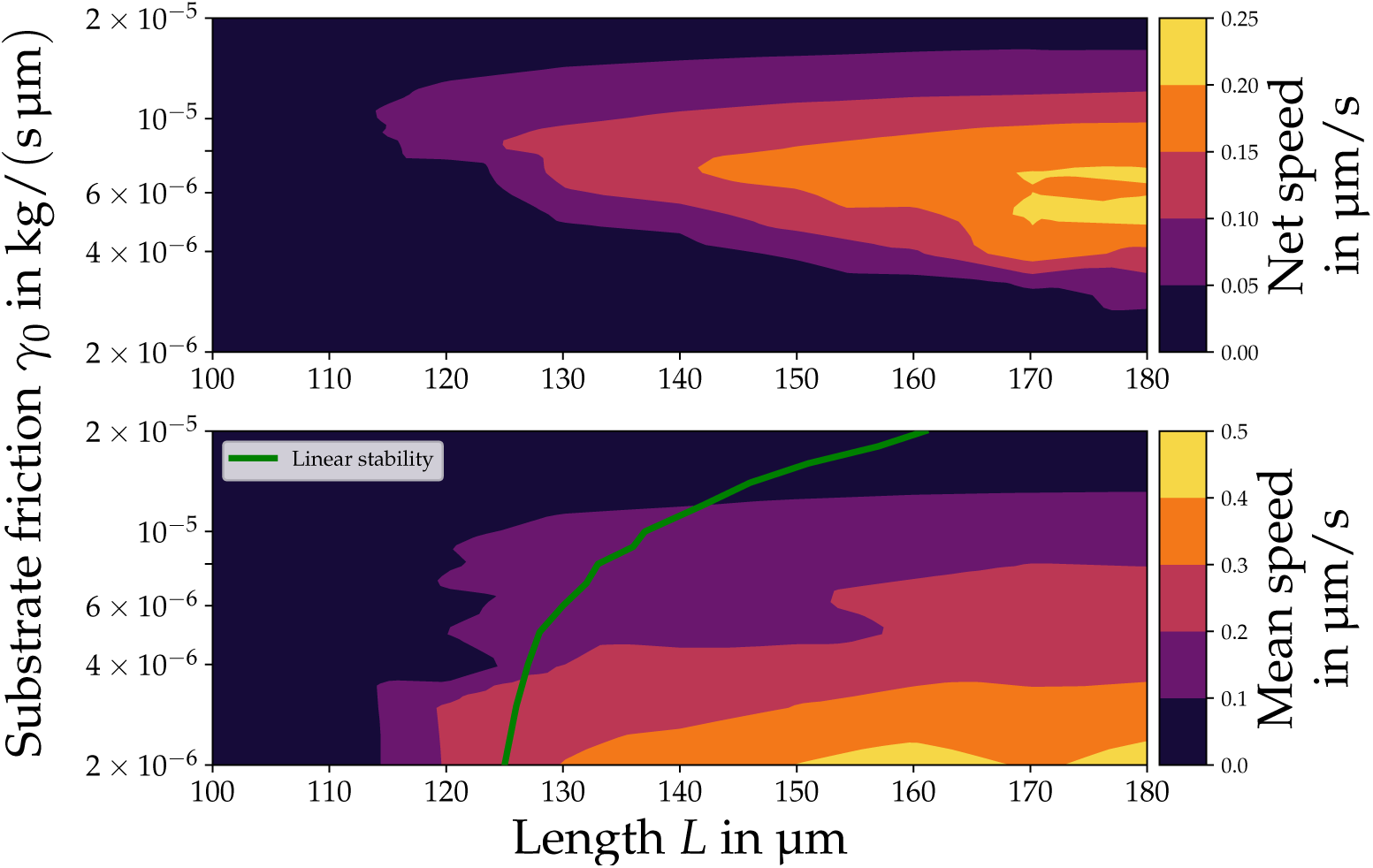
Net (top) and mean speed (bottom) for different strengths of base substrate friction *γ*_0_ and length *L*. Larger MPs possess both a higher net and mean speed. The net speed reaches a maximum for a medium base friction coefficient of *γ*_0_ = 6 × 10^−6^ kg/s while the mean speed increases with decreasing friction. There is a critical length of *L*_cr_ ≈ 120 µm in the model and *L* need to be larger than *L*_cr_ to transition from global calcium oscillations to states with motion of the boundaries. This is in qualitative agreement with the linear stability analysis that predicts the transition to an oscillatory short-wavelength instability (green line, bottom) at 130 µm for *γ*_0_ = 6 × 10^−6^ kg/s. Parameters: *B* = 3.5, *v*_slip_ = 2.5 µm/s, *α* = 0.15 and *F* = 12.3.

**Fig 6.**
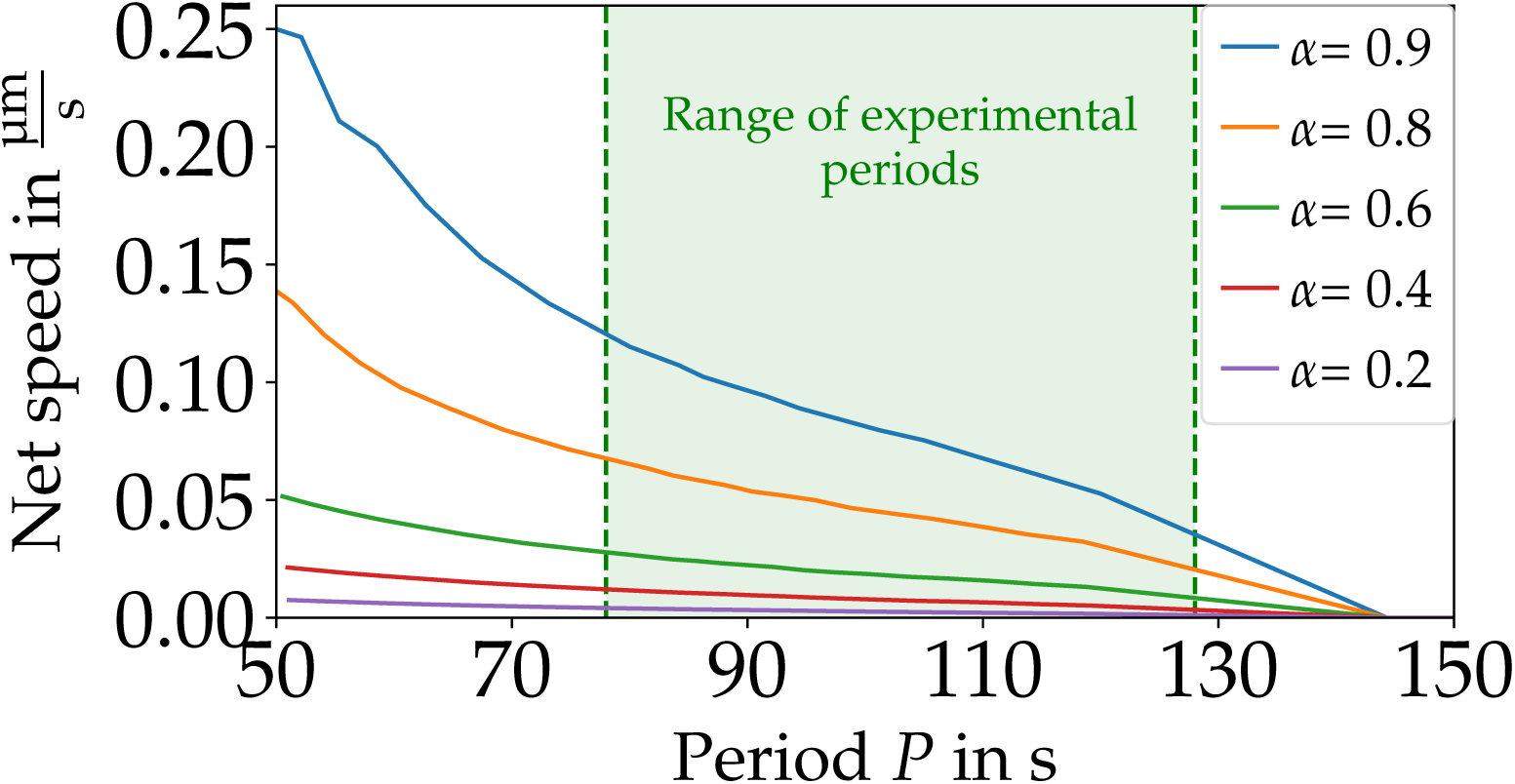
A higher period *P* of the internal dynamics results in lower net speeds for type 1 motion. For a period of more than *P* > 145 s there is a transition to global calcium oscillations without any motion. The experimentally observed periods are between 78 s and 127 s, taken from [13]. Parameters: *B* = 3.5, *v*_slip_ = 3.0 µm/s and *F* = 12.3.

Although experimental measurements of the free calcium dynamics are noisy, forward traveling calcium waves can be discerned that annihilate on collision with the front [13]. Moreover, the velocities of two different quantities can be measured: endo-and ectoplasm [13]. The ectoplasm is gel-like and connected to the MP’s membrane while the endoplasm is fluid-like. Strength and direction of the ectoplasm’s velocity correlate with measured traction forces exerted on the substrate. In the model, the gel is viscoelastic and exhibits friction with the substrate. Hence, the velocities of gel and ectoplasm should be compared. For the more common peristaltic type of MP motion, the ectoplasm’s velocity alternates periodically between forward and backward direction at a fixed position. While the velocity’s direction at front and back are equal, the center part is moving into the opposite direction with a weaker amplitude than at the boundaries. Forward ectoplasm motion has a larger amplitude than backward motion, resulting in a net speed of *v*_exp_ ≈ 0.05-0.25 µm/s with an internal period of *P*_exp_ ≈ 80 s-130 s [13, 23]. A space-time plot of the ectoplasm velocity is shown in [13] (Fig. 7a).

**Fig 7.**
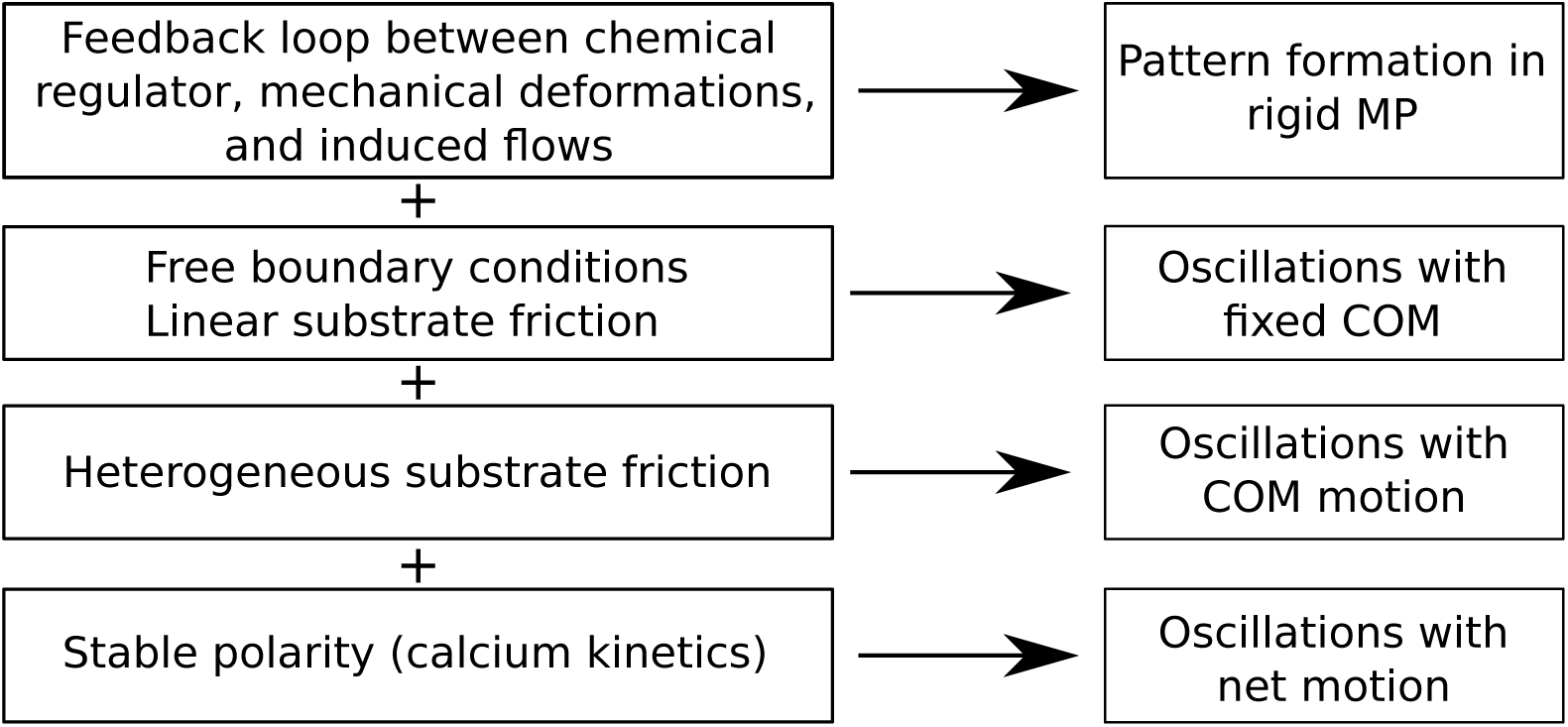
Overview of the different levels in the model and corresponding types of locomotion.

Both types of simulated movement exhibit oscillatory COM motion with net motion in each cycle. For type 1 motion (Fig. 3) the period of ≈ 95 s and net speed of 0.2 µm/s are comparable to the experiment. However, the COM is only moving while a calcium wave is propagating and is at rest otherwise which has not been observed in the experiment. Type 2 motion (Fig. 4) features continuous forward and backward motion with a net speed of 0.21 µm/s and a period of 14 s. Aside from the substantially shorter temporal period, this type does more closely resemble the experimental observations.

The calcium dynamics of both types feature traveling waves. Type 1 motion exhibit only backward traveling waves that annihilate at the rear. However, for type 2 motion waves get partly reflected and there are forward and backward traveling calcium waves. In the experiment, only forward traveling waves that annihilate at the front can be observed [13].

In addition, we can compare the experimental ectoplasm velocity with the simulated gel velocity. For type 1 motion, each calcium wave causes huge gel deformations that are accompanied by COM motion. When the calcium wave has abated, the stress in the medium relaxes and after ≈ 25 s all deformations have decayed. There is no further motion until the next calcium wave emerges. The velocity patterns in this type of motion have not been observed experimentally.

However, there are several similarities between the velocity patterns in peristaltic MP and type 2 motion. In experiment as well as simulation the gel’s velocity alternates between forward and backward direction at a given position. Usually, the gel’s velocity possesses the same direction at front and back, whereas there is opposing motion at the center. The forward velocity has a larger amplitude, which results in net motion towards the front. However, there are also differences: In the experiment, the ectoplasm’s velocity at the rear is higher than at the front which is not the case in our simulations under the chosen parameter values. As mentioned above, the period of ≈ 14 s is shorter than experimentally measured periods.

### Microplasmodia speed depends on length *L*, base friction coefficient *γ*_0_ and internal period *P*

In the parameter plane spanned by base friction strength *γ*_0_ and length *L* the type of motion as well as net and mean speed changes as displayed in Fig. 5. The mean speed is measured by the averaged speed of the boundaries. For very high values of *γ*_0_ (≥ 3 × 10^−5^ kg/s) global calcium oscillations emerge without any gel or COM motion, regardless of the length. For low values of *γ*_0_ (< 10^−6^ kg/s), there is effectively no friction between gel and substrate. In this case, the equations do not possess full rank and we omit showing these results.

In the intermediate range the length *L* strongly affects the emerging dynamics. For *γ*_0_ = 10^−5^ kg/s global calcium oscillations occur when *L* ≤ 100 µm. Increasing *L* above 110 µm gives rise to type 1 motion with a low net speed of 0.03 µm/s (Fig. 5, top). With a further increase of *L* the net speed rises up to 0.16 µm/s for *L* = 170 µm. When decreasing the base friction coefficient to *γ*_0_ = 5 × 10^−6^ kg/s, the net speed increases to 0.21 µm/s where it reaches a maximum. A further decrease yields lower net speeds. At 2 × 10^−6^ kg/s a transitions to oscillations of the COM without net motion is observed.

Additionally, we find a critical length *L*_cr_ ≈ 120 µm for the transition between global calcium oscillations and states with motion of the boundaries (Fig. 5, bottom). Our linear stability analysis predicts that there is a critical length for the transition to an oscillatory short-wavelength instability (green line). This prediction is close to the transition from global calcium oscillations to COM oscillations without net motion (bottom panel).

The existence of a critical length as well as the increasing net speed for longer MPs are qualitatively in line with experimental observations. It was found that there is a critical MP size for the onset of locomotion which is around 100 µm-200 µm [13, 61].Furthermore, previous experiments show that an increase in the longitudinal length increases the net speed [61, 62]. Note that MPs start to branch when they grow above 300 µm-500 µm [13, 21, 63] and that the model cannot capture any effects caused by this.

By changing the temporal scale *ψ* of the calcium kinetics (Eq. 5) we can adjust the period *P* of the internal dynamics for type 1 motion. In general, higher (lower) values of *ψ* result in a shorter (longer) period *P*. The net speed decreases when increasing the period *P* (Fig. 6). With a decreasing period the intervals between two calcium waves become shorter. As each wave results in net motion, the net speed decreases with higher periods until there is a transition to global calcium oscillations without any motion for periods of more than *P* ≥ 145 s. In the experiment, the maximum period observed for moving MP is 128 s, which is qualitatively in line with the maximum period of ≈ 145 s for moving MP. Consistent with our results above, a larger slip-ratio *α* increases the net speed.

To our knowledge, there is no published experimental data where the MP net speed is compared to the period of its internal dynamics. Our modeling results may hence be tested in future more detailed experimental studies of motion of MP.

## 4 Discussion

In the present work, we have extended the two-phase model with free-boundaries for poroelastic droplets from [38] in two steps. First, we introduce a nonlinear slip-stick friction between droplet and substrate. While we were able to observe back and forth motion of the boundaries in [38], we showed that COM motion is impossible with a spatially homogeneous substrate friction. The nonlinear stick-slip approach results in a heterogeneous friction coefficient and allows for COM motion. Second, we consider a nonlinear oscillatory chemical kinetics that was derived for calcium as the regulator which controls the gel’s active tension in Physarum microplasmodia (MP) [40]. It is known from previous studies [39, 40] that introducing a nonlinear calcium kinetics can lead to a uni-directional propagation of mechano-chemical waves that establishes an axis of polarity which is stable. This is in contrast to our previous work where the pure advection-diffusion dynamics of a passive regulator does not lead to the establishment of a polarity axis necessary for net motion [38].

With this extended model, we explore the conditions for self-organized motion of Physarum MP. We identified varying modes of locomotion ranging from global calcium oscillations without motion to spatio-temporal oscillations with and without net motion. Our study was restricted to one spatial dimension which can be rationalized by the experimental evidence that moving MPs have an elongated shape and motion is mostly caused by deformation waves along the longitudinal axis [62].

With the extended model, we could identify two different types of COM oscillations with net motion in addition to modes with time-periodic or irregular switching polarity. The more frequent type is characterized by mechano-chemical waves traveling from the front towards the rear that are accompanied by COM motion (Fig. 3). The second, less frequent type is characterized by mechano-chemical waves that appear alternating from front and back (Fig. 4). While both types exhibit oscillatory forward and backward motion with net motion in each cycle, in particular the trajectory and gel flow pattern of the second type resemble experimental measurements of peristaltic MP motion [13]. Interestingly, there are also experimental observations of two different types of oscillations with net motion in addition to disorganized patterns [13].

Further, we varied parameters that are accessible in experiments, such as the period of the internal dynamics *P*, the length *L*, and the base substrate friction coefficient *γ*_0_.

In the parameter plane spanned by the base friction coefficient (*γ*_0_) and the length (*L*), the model predicts that MPs need to be longer than a critical length of ≈ 120 µm to transition from global calcium oscillations to states with motion of the boundaries (Fig. 5). This result agrees with the linear stability analysis which indicated a transition to an oscillatory short-wavelength for ≈ 125 µm-130 µm. Furthermore, experimental studies found a minimal length of 100 µm-200 µm for MPs to start their motion [13, 61]. In addition, the net speed in the simulations increases with length *L* which was also observed in experiments [62].

The net speed becomes maximal for intermediate values of *γ*_0_ and larger slip-ratios *α* result in a larger net speed (Fig. 6). While the exact nature of Physarum’s friction dynamics can not be inferred from the available experimental data, introducing a nonlinear friction dynamics is crucial for COM motion. Future experiments may provide more quantitative information on the interactions between substrate and MP which could be used to improve the model.

The net speed in the simulations decreases monotonically with an increasing period (Fig. 6). For parameters leading to a period *P* > 145 s, we only found global calcium oscillations without any motion. This is qualitatively in line with experimental results, where the maximum observed period is 128 s for moving MP [13].

With the results from this work, we have been able to identify components that are essential for different types of MP locomotion. It is known from [37] that a feedback loop between a passive chemical regulator, active mechanical deformations, and induced flow is sufficient for the formation of spatio-temporal contraction patterns. In [38], we extended the model by employing free boundary conditions and linear substrate friction which yielded oscillatory and irregular motion of the MP’s boundaries with a resting COM. In the present work, we introduced nonlinear stick-slip substrate friction that enables COM motion. Moreover, inclusion of a nonlinear oscillatory calcium kinetics leads to a self-organized stable polarity which eventually allows for net motion. Fig. 7 summarizes the connection between these components and describes the corresponding types of locomotion.

Much larger forms of Physarum like mesoplasmodia and extended networks show a clear separation of gel and fluid-like phases [8, 12, 63, 64]. Recently, a study by Weber et. al [65] identified that differences in the activity between two phases can lead to a phase separation. Future extensions of our model may include this aspect by treating the phase composition as a spatially dependent local variable that includes transitions between both phases.

## Supporting information

**S1 Fig. Moving MP in body reference (top) and lab frame (bottom).** We solve our model equations in the gel’s body reference frame and the resulting quantities are defined in this frame. However, observers are located in the lab frame. The quantity’s transformation from body reference to lab frame is given by the displacement field *u* with *X*_0_ = *x*_0_ + *u*(*x*_0_). Here, *x*_0_ the position in the body reference and *X*_0_ is the position in the lab frame. Parameters from Fig. 4 in the main text.

**S2 Fig. MP net speed increases with a larger slip ratio *α*.**

**S3 Fig. Calcium concentration at the MP’s boundaries.** The magnitude of emerging waves is always higher at the MP’s front and it is moving into this direction. Parameters from Fig. 4 in the main text.

**S4 Fig. COM (orange) and boundary (blue) trajectories (top) and calcium dynamics (bottom) with irregularly switching polarity**. Parameters: *B* = 2.5, *α* = 0.1, *v*_slip_ = 3.1, *γ*_0_ = 10^−5^ kg/s, *L* = 125 µm and *F* = 15.4.

**S1 Text. Supplemental Information.** S1 Text provides the following: i) a visual comparison of a moving MP in lab and body reference frame ii) Dispersion relations for the linear stability analysis iii) A table with the default parameters.

**S1 Table.** These parameters are used throughout this work and any derivation is explicitly marked. Taken from [40].

**S1 Data.** S1 Data provides the data for all figures in the paper

## Acknowledgments

We thank Markus Radszuweit for helpful discussions about his previous work on the model, Shun Zhang and Juan Carlos del Álamo for discussions and Brian Camley for discussions. Computational resources were provided by the Institut für Theoretische Physik at the TU Berlin.

